# Emotional recognition training modifies neural response to emotional faces but does not improve mood in healthy volunteers with high levels of depressive symptoms

**DOI:** 10.1101/335042

**Authors:** Ian S. Penton-Voak, Sally Adams, Katherine S. Button, Meg Fluharty, Michael Dalili, Michael Browning, Emily Holmes, Catherine Harmer, Marcus R. Munafò

## Abstract

**IMPORTANCE:** Depression is a debilitating and highly prevalent mental health disorder. There is a need for new, effective, and scalable treatments for depression, and cognitive bias modification (CBM) of negative emotional processing biases has been suggested as one possibility. Such treatments may form the basis of ‘digital therapeutics’, that could be administered remotely and at low cost, should they prove to be effective.

**OBJECTIVES:** Study one was designed to determine neural correlates of a recently developed CBM technique for emotion recognition training; specifically, our aim was to compare the effects of training vs placebo on pre-specified regions of interest involved in emotion processing that are known to be sensitive to antidepressant treatment. Study two aimed to investigate efficacy of training on mood measures at 2 and 6-week follow-up and was powered to replicate and extend earlier findings.

**DESIGN, SETTING, AND PARTICIPANTS:** Both studies were double blind RCTs, in which participants completed five sessions of emotion recognition training or sham training, in the laboratory, over a one-week period. In study one (N=37), following this training, participants completed a novel emotion recognition task whilst undergoing fMRI. In study two (N=190), measures of mood were assessed post training, and at 2-week and 6-week follow-up. Both studies recruited analogue samples of healthy volunteers with high levels of depressive symptoms (BDI-ii > 14).

**MAIN OUTCOMES AND MEASURES:** In study one, our primary outcome was neural activation in the following pre-specified regions of interest: the bilateral amygdala, the mPFC, bilateral dlPFC, and the occipital cortex. In study two, our primary outcome was depressive symptoms over the last 2 weeks assessed using the BDI-ii at 6-week follow-up. Secondary outcomes included depressive symptoms measured using the HAM-D, and positive and negative affect assessed using the PANAS.

**RESULTS:** In both studies, CBM resulted in a change in emotion recognition bias, which (in study two) persisted for 6 weeks after the end of the CBM phase. In study one, CBM resulted in increases neural activation to happy faces compared to sad faces, with this effect driven by an increase in neural activity for happy faces. We saw this increase in activation for this contrast at both the whole brain level and among our *a priori* ROIs, specifically the mPFC and bilateral amygdala. In study two, CBM did not lead to a reduction in depressive symptoms on the BDI-ii, or on related measures of mood, motivation and persistence, or depressive interpretation bias.

**CONCLUSIONS AND RELEVANCE:** CBM of emotion recognition appears to have effects on neural activity that are similar in some respects to those induced by SSRI administration (study one), but we find no evidence that this has any effect on self-reported mood in an analogue sample of healthy volunteers with low mood (study two).

## Introduction

Mood disorders, dominated by major depression, are highly prevalent in the general population and constitute a substantial burden of disease. At present, NICE guidelines recommend psychotherapy for mild depression, and cognitive-behavioural therapy for moderate depression (National Institute for Health and Clinical Excellence 2009), but these therapies require individual intervention and therefore, while cost-effective, are expensive. This indicates a pressing need for novel approaches to improve treatments for depression, and to prevent relapse following cessation of treatment.

Understanding emotional signals is critical to successful social functioning but is disrupted in many psychiatric disorders (Cotter, Granger, Backx, Hobbs, Looi & Barnett, 2018). Negative processing biases in depression predict relapse in currently depressed and remitted patients, suggesting that these biases play a role in the onset and maintenance of depression. Neurocognitive models suggest that antidepressant medications have early effects on emotional processing that result in therapeutic benefit only after sufficient time has elapsed to allow interaction with others, where alteration in these biases gives rise to more positive social interactions (Warren, Pringle & Harmer, 2015). In support of these models, fMRI studies have provided evidence that SSRIs reduce change responses to emotional expressions, and that this change in response is associated with later improvement in mood (Warren, Pringle and Harmer, 2015).

Given the proposed causal role played by emotion processing in the onset and maintenance of depression, biases in this area may provide a potential target for behavioural, rather than pharmacological, intervention (Penton-Voak, Munaro, & Looi, 2017). We have developed a new cognitive bias modification (CBM) technique which targets the recognition of facial expression of emotions by initially assessing the threshold for detecting one emotion over another in an ambiguous expression (e.g., a blend of happiness and sadness), and then providing feedback to shift this threshold (e.g., to favour identification of happiness over sadness). Preliminary results from adults recruited from the general population indicate robust and generalizable effects on emotion perception (Griffiths et al, 2015; Dalili et al, 2016; Penton-Voak et al, 2013). A randomised controlled trial (RCT) with participants recruited from the general population on the basis of high levels of depressive symptoms on the Beck Depression Inventory ii (BDI-ii) also indicated that this intervention may have therapeutic benefit on positive affect which persists for at least two weeks (Penton-Voak et al 2012). This is consistent with recent models of the action of antidepressant medication, which suggest that drug treatment has early effects on emotional processing bias including the ability to detect positive versus negative facial expressions (Harmer, Goodwin et al. 2009). Here we investigated the neural correlates of our emotional recognition CBM intervention, and the therapeutic potential of this intervention.

Several studies show that SSRIs have a robust effect on emotion processing in the amygdala (Warren, Pringle & Harmer, 2015), an area that plays a key role in detecting the salience of emotional stimuli in the environment (Sander, Grafman, & Zalla, 2003; Santos, Mier, Kirsch, & Meyer-Lindenberg, 2011). The medial network has substantial amygdaloid and limbic connections (Price & Drevets, 2010), and altered neural activation is seen in the medial prefrontal cortex in individuals suffering from mood disorders, although the pattern of this activation varies widely between studies (Lemogne et al., 2009; Yoshimura et al., 2010, Grimm et al., 2009; Renner et al., 2015). Similarly, mood related changes in activity are found in the dorsolateral prefrontal cortex (dlPFC), a cortical area associated with the control of attention that helps regulate the amygdala through indirect inhibitory input (Davidson, 2000; Drevets, 2001). A meta-analysis of studies measuring the neural response to affective stimuli showed greater response in the amygdala, insula, and dorsal anterior cingulate cortex, and lower response in the dorsal striatum and dlPFC to negative stimuli in depressed individuals relative to healthy controls (Hamilton et al., 2012). Additionally, a review by Disner *et al.* (2011) found that biased processing of emotional stimuli in depression is associated with greater amygdala reactivity, as well as left dlPFC hypoactivity and right dlPFC hyperactivity.

Study 1 aimed to identify changes in the neural correlates of emotion recognition following CBM, by administering five days of the emotion recognition training intervention (or a sham training procedure) and then scanning participants using fMRI while performing a face perception task that has been previously used to investigate the effects of SSRIs on the processing of emotion facial expressions (Godlewska et al 2012). We hypothesised that emotional recognition training would reduce amygdala responses to negative facial expressions. We also hypothesised that training would alter activity in a) the occipital cortex, as it is highly connected to the amygdala and is sensitive to attentional change in response to emotional stimuli, and b) the prefrontal cortex, which exerts effects on circuitry implicated in pharmacological and psychological treatment for depression. Based on previous findings, we established the following areas as our regions of interest (ROIs) for comparing neural activation in individuals suffering from low mood in our intervention and control conditions: the bilateral amygdala, the mPFC, bilateral dlPFC, and the occipital cortex.

Study 2 was a RCT using an analogue sample of participants recruited from the general population on the basis of high levels of depressive symptoms on the Beck Depression Inventory ii (BDI-ii), in a direct replication of earlier work (Penton-Voak et al 2012) using a larger sample with long-term follow-up. The CBM procedure was identical to Study 1 – participants were randomised to receive either five days of the emotion recognition training intervention, or a sham training procedure. Participants completed a series of assessments of mood and anxiety at 2-week and 6-week follow-up after the end of treatment. We hypothesised that participants randomised to the emotion recognition training intervention would reduce lower symptoms of depression on the BDI-ii over the previous two weeks at 6-week follow-up (our primary outcome).

## Study One

### Methods

#### Participants

We recruited adults from the staff and students at the University of Bristol and from the general population who reported depressive symptoms (defined as a score of 14 or more on the BDI-ii) (Beck, Steer, & Brown, 1996). Participants were recruited via email lists and local advertisements.

Upon arrival, participants provided informed consent and inclusion/exclusion criteria were assessed. Screening consisted of structured clinical interview for DSM-IV: Clinical Interview Schedule; CIS-R (Lewis, Pelosi, Araya, & Dunn, 1992), the Altman Self-Rating Mania Scale; ASRM (for bipolar disorder) (Altman, Hedeker, Peterson, & Davis, 1997) and medical history. After initial screening we also collected baseline data on age, sex, ethnicity, alcohol, tobacco and caffeine use, previous history of depression (treated and non-treated), intelligence (National Adult Reading Test, NART) (Nelson, 1982), number of years of education, social network size (SNS), and current and past history of psychiatric treatment. Criteria for exclusion were a diagnosis of primary anxiety disorder, psychosis, bipolar disorder, or substance dependence (other than nicotine and caffeine) as defined by DSM-IV; current use of an illicit drug (except cannabis); being at clinically significant risk for suicidal behaviour; use of psychotropic medication in the last 5 weeks prior to the study; major somatic or neurological disorders and concurrent medication that could alter emotional processing (including active treatment with counseling, cognitive behavioural therapy, or other psychotherapies).

The study was approved by the Faculty of Science Research Ethics Committee at the University of Bristol. On completion of the final study session, participants were reimbursed £60 for their time and expenses.

#### Study design and intervention

We used minimization to allocate participants to either a training procedure designed to promote the perception of happiness over sadness in ambiguous emotional expressions, or a control procedure designed to elicit no change in perception of emotional expression, in order to ensure the groups were balanced for baseline BDI-ii symptoms (grouped according to a score of 14-19, or 20+). This was done by an experimental collaborator at the Bristol Randomised Trials Collaboration, and testing was double-blind. Participants were randomised to either a training procedure designed to promote the perception of happiness over sadness in ambiguous emotional expressions, or a control procedure designed to elicit no change in perception of emotional expression. In brief, the CBM intervention consists of three phases. First, in the baseline phase, images from a 15 face morph sequence that runs from happy to sad facial expressions are presented one at a time, with participants asked to judge whether the face is happy or sad. This allows the ‘balance point’ at which participants shift from a ‘happy’ judgement to a ‘sad’ judgement to be calculated – a measure of cognitive bias. In the training phase, feedback (correct/incorrect) is used to shift the participant’s balance point. In the training condition, the ‘correct’ classification is shifted towards ‘happy’; the two images nearest the balance point that the participant would have previously classified as ‘sad’ at baseline are considered ‘happy’ in terms of providing feedback. Feedback in the control condition is based directly on baseline performance, and has no effect on responses. Sessions last 20 minutes and are fully automated. Methods are described in detail elsewhere (Penton-Voak et al 2012, 2013). Participants completed computerised training or control procedures once a day over five consecutive days (Monday to Friday). fMRI acquisition took place after the completion of training during the last session.

#### Mood Assessment

Mood assessments via questionnaire measures were taken at baseline and at the end-of-treatment. End-of-treatment follow-up included a visual analogue scale rating of how friendly the participant thought the experimenter was, to ensure that there were no differences between treatment conditions that may have affected blinding. The questionnaire measures included the Beck Depression Inventory (BDI-ii) (Beck, 1996), the Beck Anxiety Inventory (BAI) (Beck, 1988), the Hamilton Rating Scale for Depression (HAM-D) (Hamilton, 1960), and the Positive and Negative Affect Schedule (PANAS) (Watson, Clark, & Tellegen, 1988).

#### Functional MRI Data Acquisition

Image data were acquired with a Siemens Skyra 3 tesla MR system with a 32-channel head coil. During functional imaging, echo planar T2*-weighted images (EPIs) were acquired in a transversal direction parallel to the AC-PC line (36 slices, repetition time (TR) = 2,520 ms, echo time (TE) = 30 ms, flip angle = 90°, field of view (FOV) = 192 mm^2^, imaging matrix = 64 × 64, slice thickness = 3 mm, slice gap = 0.8 mm, voxel size = 34 mm^3^). After the main experimental task, a three-dimensional T1-weighted Magnetization-Prepared Rapid Gradient Echo (MPRAGE) image volume was acquired as anatomical reference (192 slices, TR = 1,800 ms, TI = 800 ms, TE = 2.25 ms, imaging matrix = 256 × 256, FOV = 240 mm^2^, flip angle = 9°, slice thickness = 0.9 mm, and voxel size = 0.8 mm^3^).

#### Functional MRI Behavioural Task

During fMRI scanning, participants completed a simple sex discrimination task involving the rapid presentation of sad, happy, and fearful facial expressions. In this task, thirteen 30 sec blocks of a baseline fixation cross (condition A) were interleaved with twelve 30 sec blocks of the emotional task – four blocks of sad (condition B), four blocks of happy (condition C), and four blocks of fear (condition D). During each emotional block participants viewed 10 emotional faces (5 female) all derived from a standard set of pictures of facial affect (Tottenham et al, 2009). Each face was presented for 150 ms and participants were asked to report the sex of the face via button press using an MRI-compatible keypad. The within block interstimulus intervals (ISI) was set at 2900 ms. During the course of the 8.5 min experiment, participants completed the blocks in the following order: ADACABADACABADACABADACABA. Stimuli were presented on a personal computer using E-Prime (version 1.0; Psychology Software Tools Inc., Pittsburgh, PA, USA) and a cloned projection displayed to participants on an opaque screen located at the head of the scanner bore, which participants viewed using angled mirrors. Stimulus presentation and participant button presses were registered and time-locked to fMRI data using E-Prime. Both accuracy (correct sex discrimination) and reaction times were recorded.

#### Functional MRI Pre-Processing and Statistical Analysis

fMRI data processing was carried out using FEAT (FMRI Expert Analysis Tool) Version 6.00, part of FSL (FMRIB’s Software Library, www.fmrib.ox.ac.uk/fsl). Motion correction was applied using a rigid body registration to the central volume. Gaussian spatial smoothing was applied with a full width half maximum of 5 mm. High pass temporal filtering was applied using a Gaussian-weighted running lines filter, with a 3 dB cut-off of 120 sec. We fitted a general linear model with three explanatory variables: ‘sad faces’, ‘happy faces’ and ‘fearful faces’. All explanatory variables were convolved with a default haemodynamic response function (Gamma function, delay = 6 sec, standard deviation = 3 sec), and filtered by the same high pass filter as the data. The full model was simultaneously regressed to the data, giving the best-fitting amplitudes for each explanatory variable. This first-level analysis yielded beta difference images (contrast of parameter estimates; COPEs) for the blood oxygen level dependent (BOLD) response for happy>rest, sad>rest, and fearful>rest, as well as happy>sad, happy>fear, happy>sad+fear, and sad>fear.

Functional imaging data was transformed to standard space using each participant’s high resolution structural image (i.e., their MPRAGE scan). First, each participant’s functional images were registered to their structural scan using boundary-based registration (BBR, Greve & Fischl, 2009). Second, the structural brain volume was separated into brain and non-brain tissue using voxel-based morphometry (VBM8) in Statistical Parametric Mapping (SPM8, Wellcome Department of Cognitive Neurology, London, UK, http://www.fil.ion.ucl.ac.uk/spm) software. Third, the resultant brain extracted structural images were registered to the Montreal Neurological Institute (MNI) template brain, using a combination of linear (FLIRT, Jenkinson, Bannister, Brady, & Smith, 2002) and non-linear (FNIRT) transformations. Finally, the different transformation matrices and non-linear warp fields were combined to allow transformation from functional to “standard” template space for the purpose of group analysis.

At the group level, individual participant’s first-level difference maps were compared using permutation testing in FSL (RANDOMISE: FSL’s tool for nonparametric permutation inference) within our regions of interest: the bilateral amygdala, bilateral dorsolateral prefrontal cortex (dlPFC), occipital cortex, and medial prefrontal cortex (mPFC). RANDOMISE allows spatial interrogation of effects within the ROI for pair-wise comparisons of lower-level contrasts. Furthermore, RANDOMISE is preferable to FLAME (FMRIB’s Local Analysis of Mixed Effects, Beckmann, Jenkinson, & Smith, 2003; Woolrich, Behrens, Beckmann, Jenkinson, & Smith, 2004; Woolrich et al., 2009) for small volumes, such as ROIs. As RANDOMISE can use thresholded masks and makes voxel-wise comparisons, it has better spatial precision for measuring neural activation in a particular region. This is especially advantageous in regions where measuring activation has proven difficult, such as the amygdala, where measurements of activation have been found to be confounded by an adjacent large vein draining distant brain regions (Boubela et al., 2015). We applied a cluster-based correction for multiple comparisons across the voxels in the ROIs using a threshold of Z =2.3 and a corrected cluster level P < .05.

As emotion recognition training trained individuals to identify ambiguous expressions as happy rather than sad, our main contrast of interest was therefore happy>sad. We examined happy>fear and happy>sad+fear to explore whether effects generalised to other negative emotions. We also examined the three “emotion” > rest contrasts to explore which emotions underpinned any observed effects. Where group differences for emotion contrasts were significant, mean percent signal change values were extracted for each participant and compared across conditions to characterise the specific effect.

We derived masks for left and right amygdala from the Harvard-Oxford subcortical anatomical atlas, selecting voxels with a > 50% probability of lying within the amygdala. We used a derived mask for the occipital cortex from the Harvard-Oxford cortical anatomical atlas occipital lope, using a probability threshold of 50%. For the MPFC and DLPFC we downloaded the reverse-inference activation masks (searched terms MPFC and DLPFC respectively) from http://neurosynth.org. For the DLPFC we then restricted the Neuro-synth mask to the medial frontal gyrus, manually removed voxels unattached to main cluster and then applied a dilation (fslmaths) to generate smooth masks for right (central coordinates 25, 79, 50) and left DLPFC (central coordinates 66, 78, 51). A similar procedure was used for the MPFC mask (central coordinates 45, 89, 41). Where group differences for emotion contrasts were identified, mean percent signal change values were extracted for each participant and compared across groups to characterise the specific effect.

Whole brain higher-level analysis was carried out using FLAME (FMRIB’s Local Analysis of Mixed Effects (Beckman et al, 2003, Woolwich et al, 2004). Activations were identified using cluster-based thresholding of statistical images with a height threshold of Z >

2.3 and a (whole-brain corrected) spatial extent threshold of P < 0.05. We used the Harvard-Oxford Cortical and Sub-Cortical Structural Atlases and an automatic atlas query tool ‘autoaq’ to determine in which structures any activation resided.

### Results

#### Characteristics of Participants

A total of 36 participants (24 female) aged 18 to 33 years (*M* = 22, *SD* = 4) were recruited. Due to a randomisation error, there were 19 participants in the intervention condition and 17 participants in the control condition. All participants were right-handed (EHI score ≥ 50). The characteristics of participants by condition are shown in Table 1.

**Table 1.** **Characteristics of Participants (Studies 1 and 2)**

|  | Study 1 |  | Study 2 |  |
| --- | --- | --- | --- | --- |
|  | Intervention (*n* = 19) | Control (*n* = 17) | Intervention (*n* = 95) | Control (*n* = 95) |
| Age | 21 (4) | 23 (4) | 22 (4) | 22 (5) |
| Sex (female) | 13 (68%) | 11 (65%) | 69 (73%) | 69 (73%) |
| Ancestry (European) |  |  | 69 (73%) | 64 (67%) |
| NART Score | 36.47 (6.70) | 33.00 (8.73) | 38.27 (7.08) | 38.26 (6.86) |
| Years of Education | 15.18 (1.63) | 16.53 (2.85) | 15.57 (2.43) | 15.90 (2.44) |
| CISR Score | 16.84 (9.34) | 15.06 (8.56) | 17.71 (9.79) | 16.78 (11.14) |
| ASRM Score | 3.42 (2.39) | 3.00 (1.66) | 2.88 (2.17) | 3.00 (2.39) |
| BDI-ii Screening | 25.21 (8.50) | 24.12 (6.75) | 25.00 (7.48) | 24.55 (8.72) |
| BDI-ii Baseline | 19.00 (9.10) | 18.18 (6.57) | 21.05 (9.95) | 20.93 (10.13) |
| BDI-ii End-of-Treatment | 14.37 (5.73) | 16.41 (6.96) | 17.63 (9.81) | 16.98 (10.71) |
| BDI-ii Follow-Up (2-week) | n/a | n/a | 16.15 (9.81) | 15.73 (10.99) |
| BDI-ii Follow-Up (6-week) | n/a | n/a | 13.17 (9.62) | 14.01 (10.23) |
| BAI Total Baseline | 12.95 (8.20) | 14.94 (8.54) | 14.60 (8.85) | 15.86 (10.39) |
| BAI Total End-of-Treatment | 9.95 (6.70) | 12.71 (10.80) | 11.30 (7.96) | 11.07 (9.05) |
| BAI Follow-Up (2-week) | n/a | n/a | 10.83 (9.83) | 10.34 (9.60) |
| BAI Follow-Up (6-week) | n/a | n/a | 10.33 (9.34) | 10.15 (9.05) |
| HAM-D Total Baseline | 15.05 (5.34) | 15.47 (5.43) | 13.25 (5.68) | 13.41 (6.38) |
| HAM-D Total End-of-Treatment | 11.74 (5.63) | 14.81 (5.12) | 9.13 (5.12) | 8.90 (5.58) |
| HAM-D Follow-Up (2-week) | n/a | n/a | 9.43 (5.76) | 9.53 (6.53) |
| HAM-D Follow-Up (6-week) | n/a | n/a | 8.08 (5.45) | 9.06 (6.06) |
| PANAS Positive Score Baseline | 17.26 (6.45) | 17.59 (4.35) | 16.91 (5.33) | 18.06 (7.33) |
| PANAS Positive Score End-of-Treatment | 18.05 (7.15) | 19.41 (5.81) | 17.81 (6.55) | 18.69 (7.36) |
| PANAS Positive Follow-Up (2-week) | n/a | n/a | 18.30 (6.78) | 19.69 (7.75) |
| PANAS Positive Follow-Up (6-week) | n/a | n/a | 19.80 (8.36) | 19.99 (7.90) |
| PANAS Negative Score Baseline | 15.53 (5.38) | 16.94 (5.87) | 15.84 (5.55) | 15.77 (6.06) |
| PANAS Negative Score End-of-Treatment | 13.53 (3.39) | 16.65 (6.08) | 15.10 (4.82) | 14.54 (5.23) |
| PANAS Negative Follow-Up (2-week) | n/a | n/a | 14.76 (5.14) | 15.20 (6.15) |
| PANAS Negative Follow-Up (6-week) | n/a | n/a | 14.34 (4.47) | 14.31 (5.17) |
| Experimenter Friendliness | 8.73 (1.59) | 8.69 (1.52) | 8.29 (1.66) | 8.48 (1.65) |

Values represent mean (standard deviation) for continuous variables, and number (percentage) for categorical variables.

#### Behavioural Results

Participants in the intervention condition showed a shift in balance point compared to participants in the control condition after 5 sessions, adjusting for their session 1 baseline balance point (adjusted mean difference 4.65, 95% CI = 2.95 to 6.36, P < .001). Mean balance points at baseline and test for intervention and control conditions are presented in Figure 1. A mixed model ANOVA of questionnaire score data with a between-subjects factor of training condition (intervention, control) and within-subjects factor of time (baseline, follow-up) indicated evidence of main effect of time across measures (*F*s [1, 33] = 6.66 to 9.59, Ps ≤ .014), reflecting an improvement of mood from baseline to follow-up, except for the PANAS positive and negative scores (*F*s [1, 33] = 2.08 to 3.06, Ps ≥ .089), where the direction of effect was consistent with other measures but the statistical evidence weaker. We did not find clear evidence of a main effect of training condition across all measures (*F*s [1, 33] = 0.07 to 2.72, Ps ≥ .10), or any clear evidence of an interaction between time and training condition across measures (*F*s [1, 33] = 0.24 to 2.68, Ps ≥ .11). Due to a programming error, data from the sex discrimination task were not recorded.

**Figure 1.**
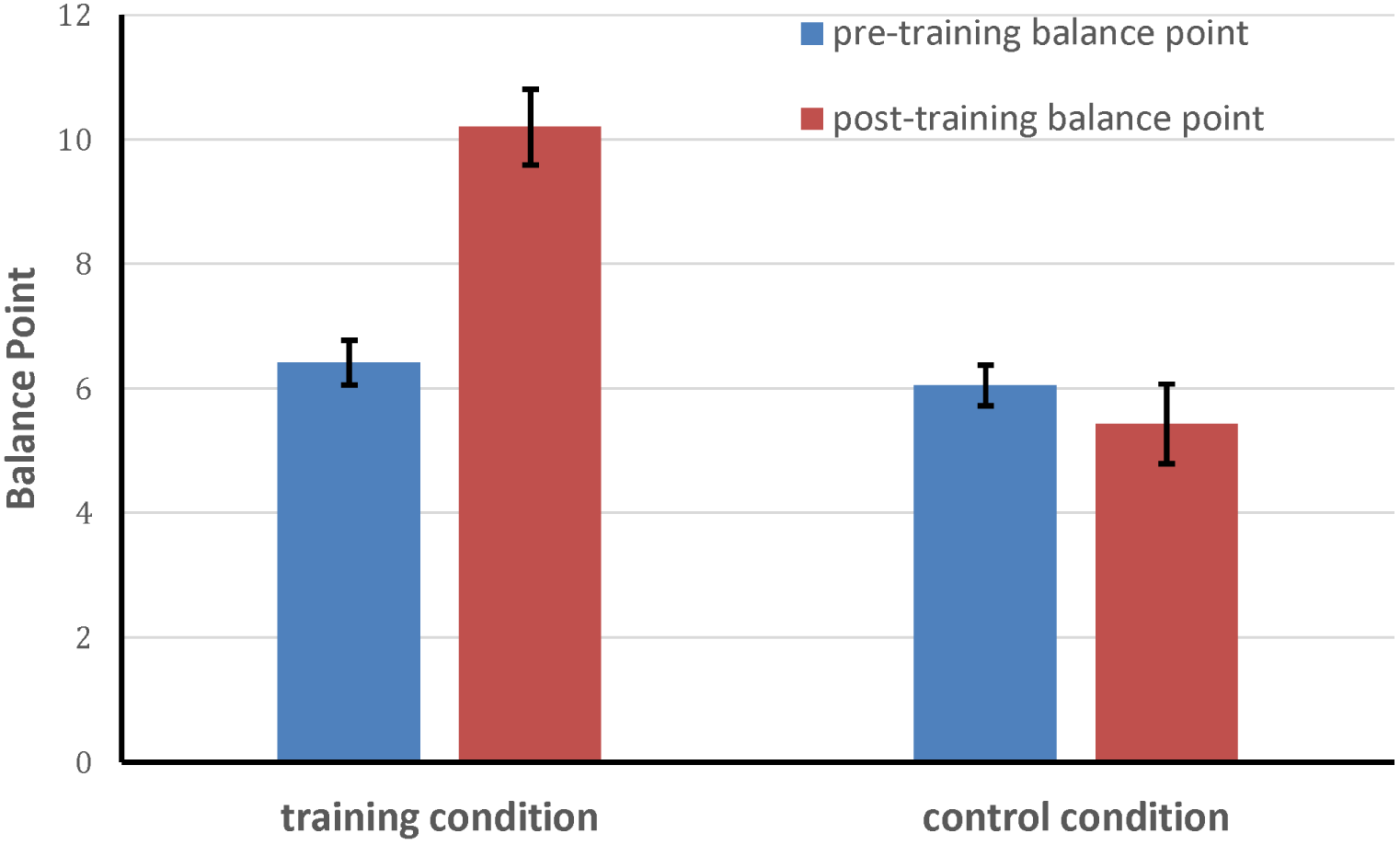
**Mean (+/- 1SE) balance points pre‐ and post-training (session 5) for training and control groups. Higher balance points indicate a bias towards happy responses.**

#### Functional MRI Results (regions of interest)

Due to a lost imaging data file, we analysed the fMRI data of 35 participants (19 intervention, 16 control). Our ROI analyses showed evidence of increased activation to the happy>sad contrast in the intervention condition relative to the control condition, but only in the left, and not the right, amygdala (FWE corrected P < .05, central coordinates 57, 61, 27; see Figure 2). There were no group differences on the happy>sad contrast in the other ROIs (occipital cortex, dlPFC, or mPFC).

**Figure 2.**
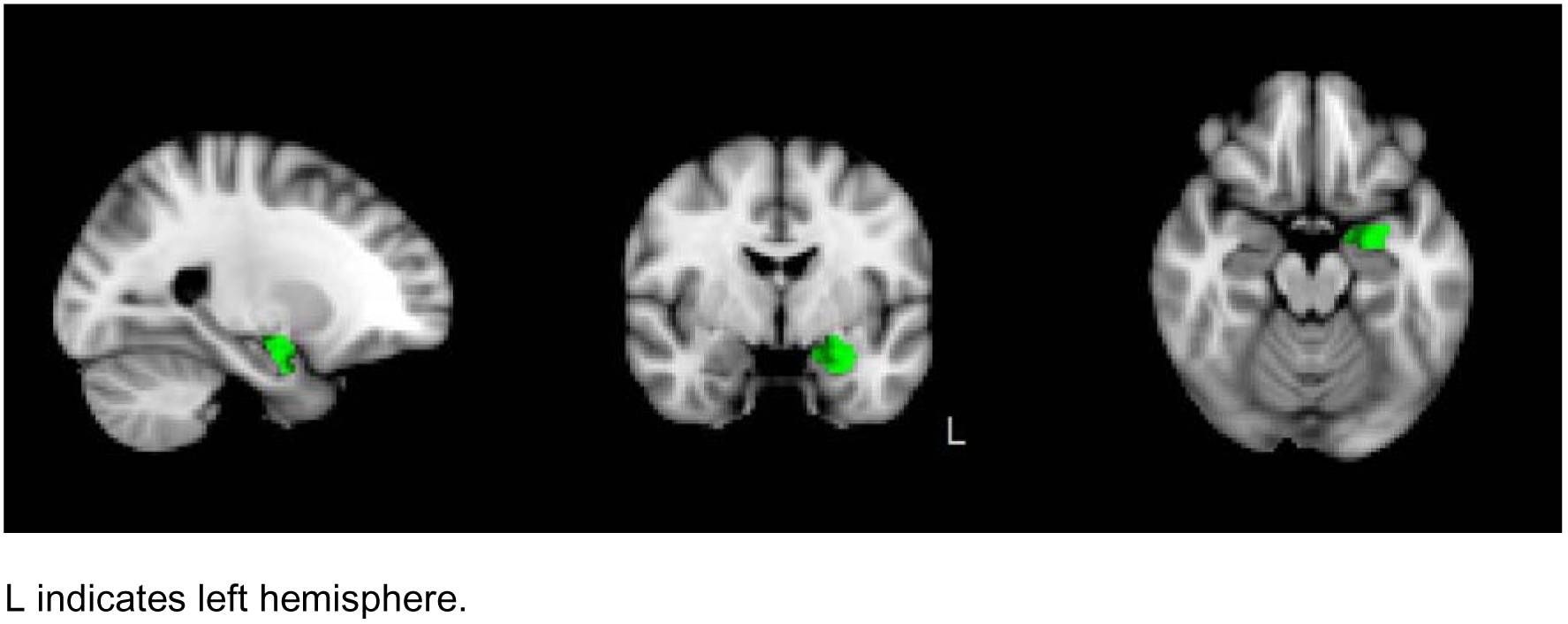
**Increased activity for the happy>sad contrast in the left amygdala for the intervention condition relative to the control condition (cluster corrected p < .05).**

Training also increased BOLD activation to happy>fear and happy>sad+fear contrasts in the left amygdala. These group differences were driven by increased BOLD activation to happy faces in the intervention condition compared to the control condition, with higher BOLD activation to the happy>rest contrast in both the left and right amygdala and also in the mPFC (see Figure 3). Percent signal change in activation for happy faces relative to rest for both the intervention and control conditions in the bilateral amygdala and mPFC is shown in Figure 4. There were no group differences for sad>rest, fear>rest, or sad>fear, and no evidence for increased activation in any contrasts for the control condition relative to the intervention condition. There was no evidence for group differences on any contrasts in any of the other ROIs.

**Figure 3.**
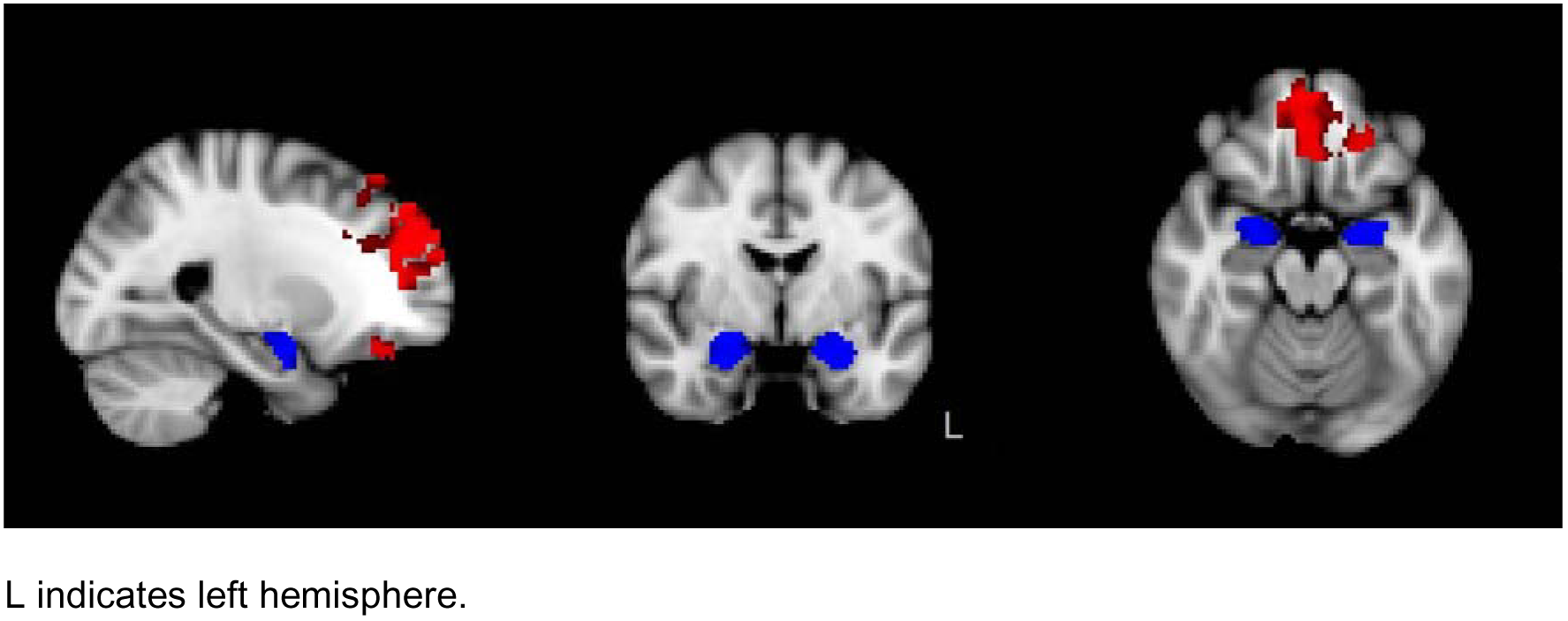
**Increased activity for the happy>rest contrast in the bilateral amygdala and mPFC for the intervention condition relative to the control condition (cluster corrected P < .05).**

**Figure 4.**
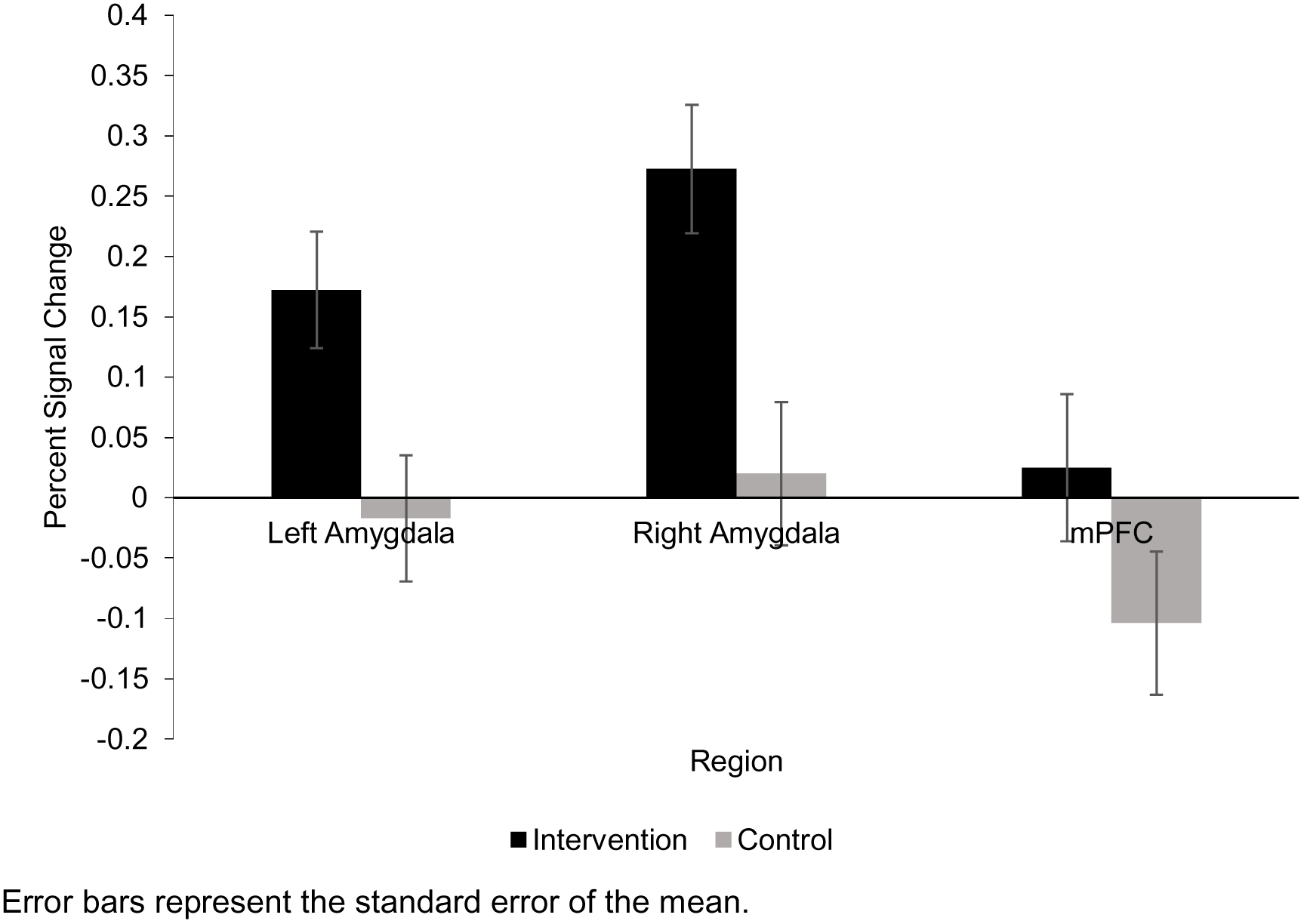
**Percent signal change of activation for the happy>rest contrast in the left amygdala, right amygdala, and mPFC for the intervention and control conditions.**

To characterise the specific effect of training in the left and right amygdala between conditions for each of our three “emotion” > rest contrasts, we conducted a post-hoc repeated measures mixed model ANOVA of percent signal change with a between-subjects factor of training condition (intervention or control) and within-subjects factors of hemisphere (left or right) and emotion (happy, sad, or fear). We observed evidence of a main effect of training condition (*F* [1, 33] = 6.53, P = .015), where participants in the intervention condition showed greater activation across contrasts relative to the control group. We also found evidence of a main effect of hemisphere (*F* [1, 33] = 12.10, P = .001), where participants showed greater activation in the right amygdala compared to the left amygdala. We did not find clear evidence for any interactions between factors (P s > .22). Activation for each condition by contrast and hemisphere is shown in Figure 5. Independent samples t-tests indicated evidence of greater activation for the intervention condition relative to the control condition for the happy>rest contrast in both the left (mean difference = 0.189, 95% CI 0.044 to 0.334, P = .012) and right (mean difference = 0.253, 95% CI 0.069 to 0.436, P = .008) amygdala. We also found evidence of greater activation for the intervention condition relative to the control condition for the fear>rest contrast in the right amygdala (mean difference = 0.148, 95% CI 0.010 to 0.286, P = .036).

**Figure 5.**
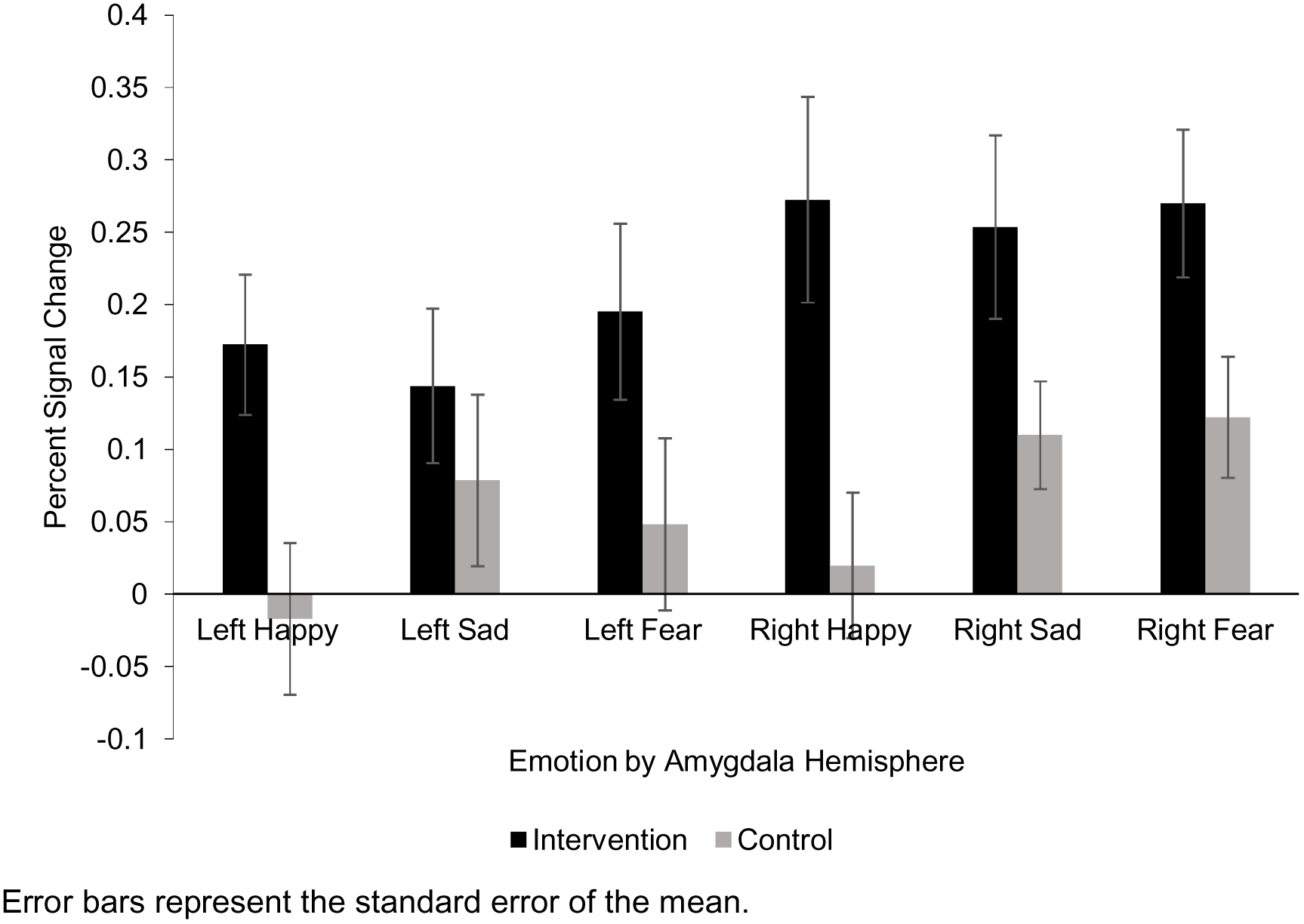
**Percent signal change of activation for each “emotion”>rest contrast in the left and right amygdala for the intervention and control conditions.**

#### Functional MRI results (whole brain)

The higher-level whole brain analysis indicated evidence of main effects of training condition on the happy≥sad+fear contrast, with participants in the intervention condition showing greater activation than participants in the control condition. The Harvard-Oxford Sub-Cortical Structural Atlas and an automatic atlas query tool ‘autoaq’ (http://brainder.org/2012/07/30/automatic-atlas-queries-in-fsl) indicated a cluster of activation (402 voxels) encompassing the brainstem, peak Z-score 3.50 (2, −24, −20) and a second cluster encompassing the left amygdala (our a priori ROI), 448 voxels, peak Z-score 4.19 (−36, 0, −20) (see Figure 6). We also explored the other contrasts and found evidence of greater activation in the intervention condition for happy>fear and happy>rest (see Figure 7). No other group differences were observed. Full results are reported in Table 2.

**Figure 6.**
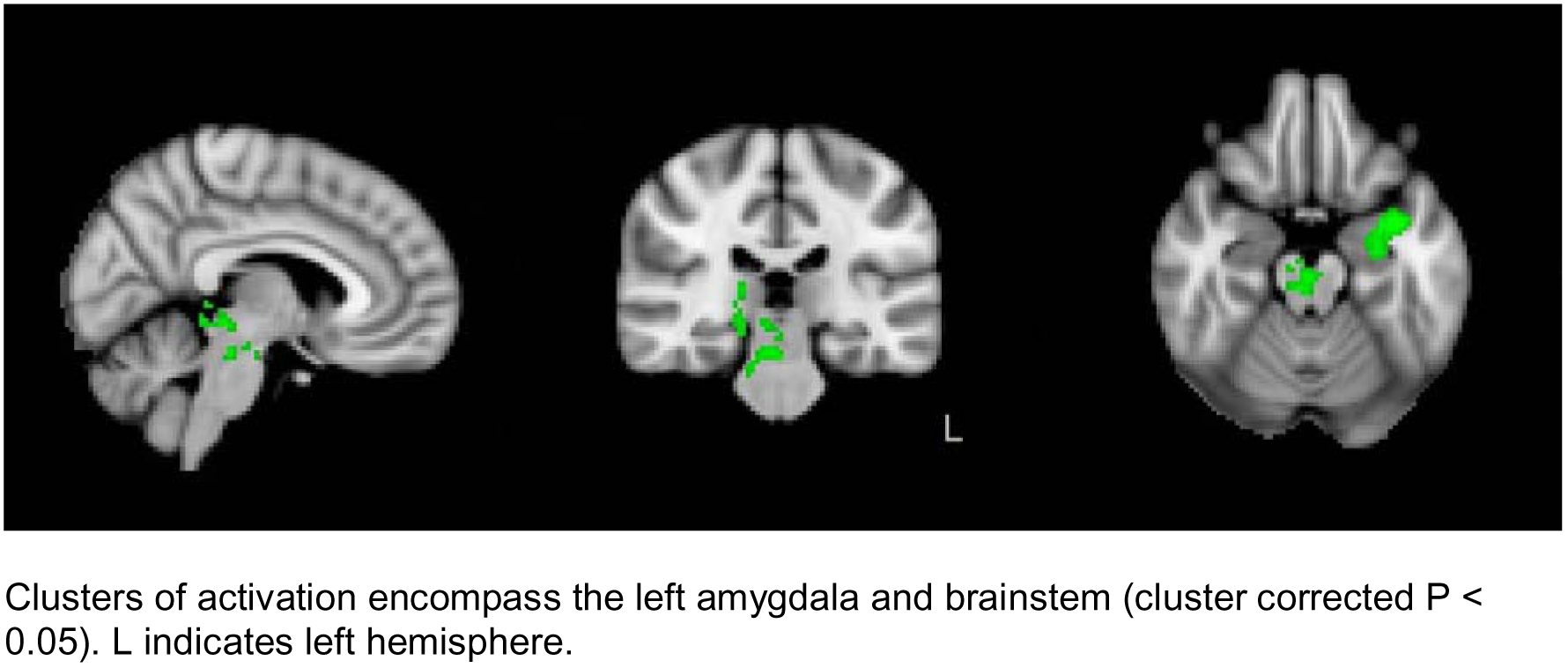
**Increased activity observed in whole brain analysis for the happy>sad+fear contrast for the intervention condition relative to the control condition.**

**Figure 7.**
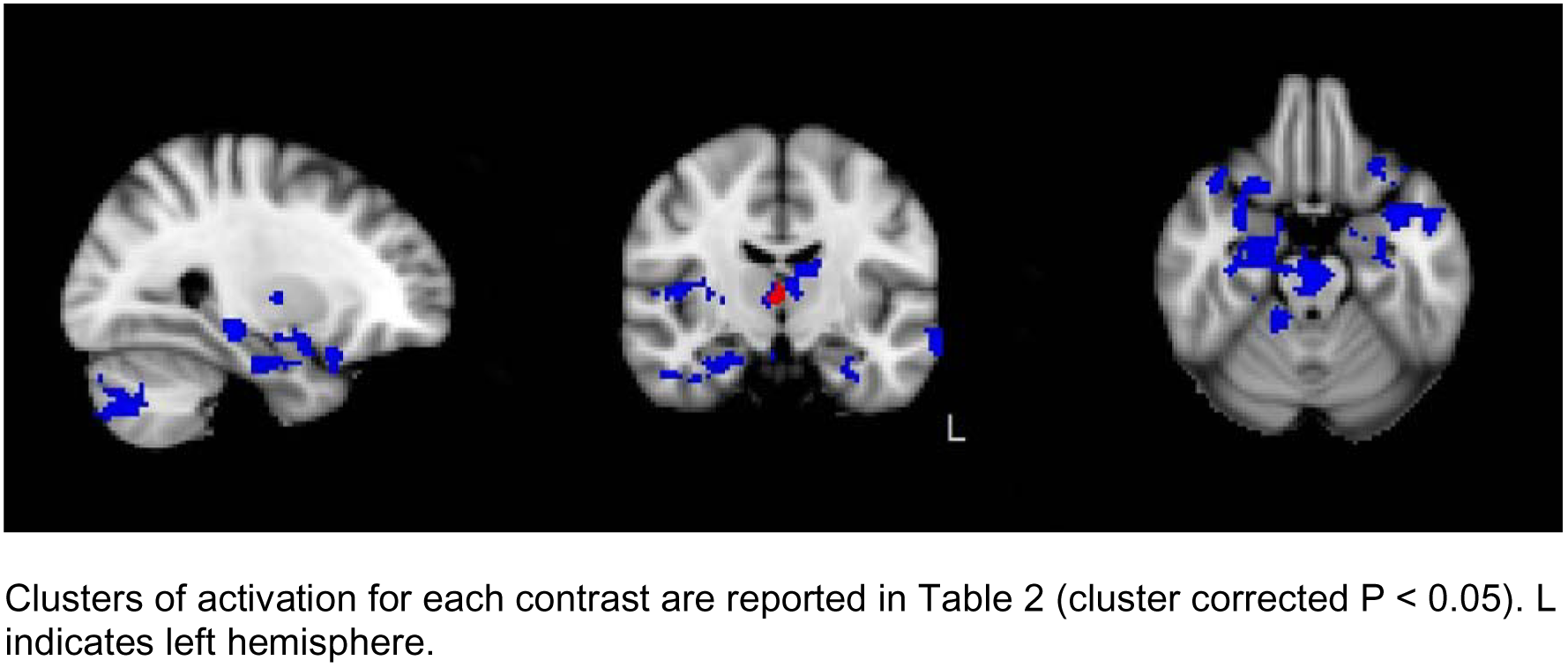
**Increased activity observed in whole brain analysis for the happy>fear (red) and happy>rest (blue) contrasts for the intervention condition relative to the control condition.**

**Table 2.**
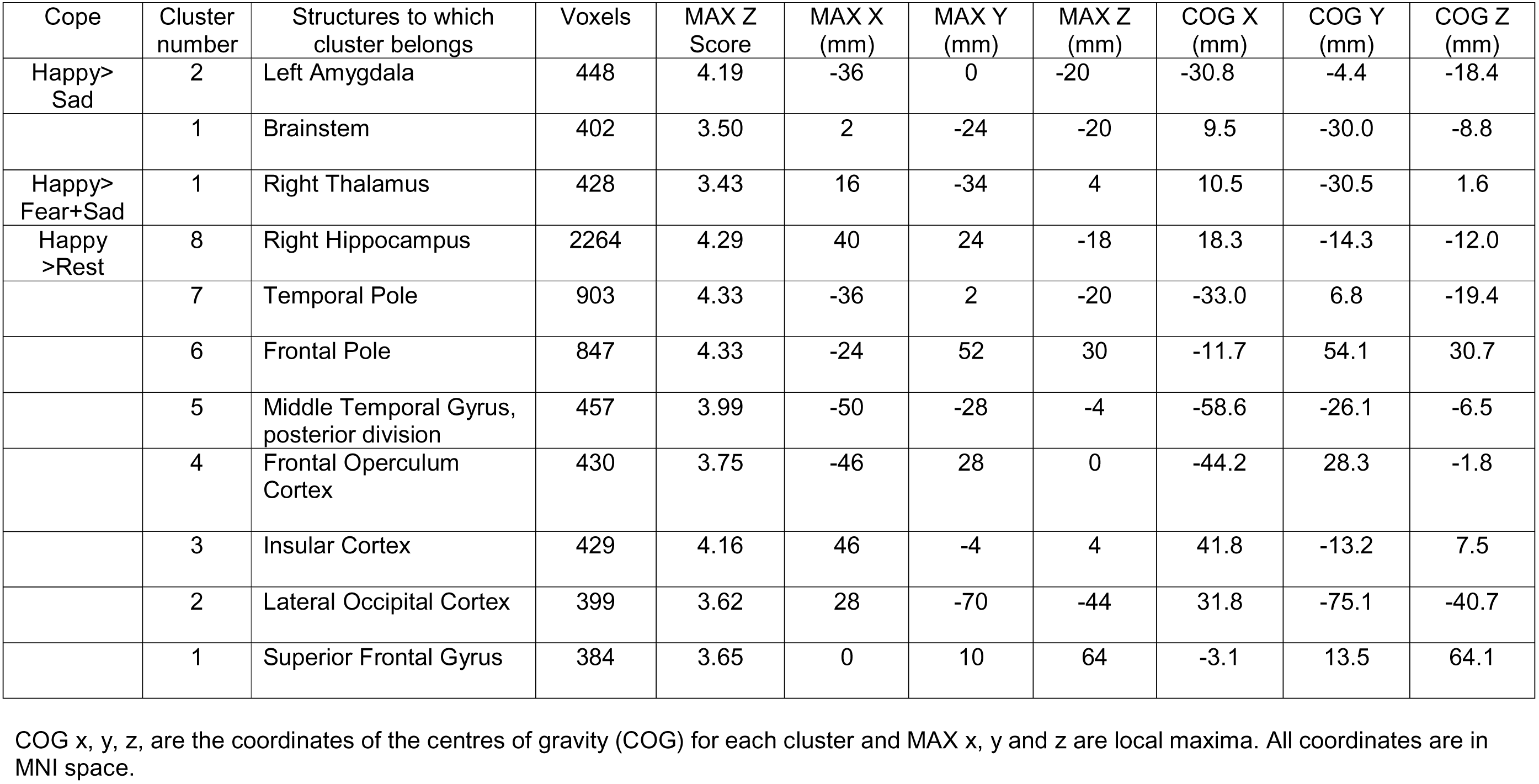
**Summary of Results of Whole Brain Analysis.**

| Cope | Cluster number | Structures to which cluster belongs | Voxels | MAX Z Score | MAX X (mm) | MAX Y (mm) | MAX Z (mm) | COG X (mm) | COG Y (mm) | COG Z (mm) |
| --- | --- | --- | --- | --- | --- | --- | --- | --- | --- | --- |
| Happy>Sad | 2 | Left Amygdala | 448 | 4.19 | -36 | 0 | -20 | -30.8 | -4.4 | -18.4 |
|  | 1 | Brainstem | 402 | 3.50 | 2 | -24 | -20 | 9.5 | -30.0 | -8.8 |
| Happy>Fear+Sad | 1 | Right Thalamus | 428 | 3.43 | 16 | -34 | 4 | 10.5 | -30.5 | 1.6 |
| Happy>Rest | 8 | Right Hippocampus | 2264 | 4.29 | 40 | 24 | -18 | 18.3 | -14.3 | -12.0 |
|  | 7 | Temporal Pole | 903 | 4.33 | -36 | 2 | -20 | -33.0 | 6.8 | -19.4 |
|  | 6 | Frontal Pole | 847 | 4.33 | -24 | 52 | 30 | -11.7 | 54.1 | 30.7 |
|  | 5 | Middle Temporal Gyrus, posterior division | 457 | 3.99 | -50 | -28 | -4 | -58.6 | -26.1 | -6.5 |
|  | 4 | Frontal Operculum Cortex | 430 | 3.75 | -46 | 28 | 0 | -44.2 | 28.3 | -1.8 |
|  | 3 | Insular Cortex | 429 | 4.16 | 46 | -4 | 4 | 41.8 | -13.2 | 7.5 |
|  | 2 | Lateral Occipital Cortex | 399 | 3.62 | 28 | -70 | -44 | 31.8 | -75.1 | -40.7 |
|  | 1 | Superior Frontal Gyrus | 384 | 3.65 | 0 | 10 | 64 | -3.1 | 13.5 | 64.1 |

COG x, y, z, are the coordinates of the centres of gravity (COG) for each cluster and MAX x, y and z are local maxima. All coordinates are in MNI space.

### Discussion

Our results suggest that emotion recognition training increases neural activation to happy faces compared to sad faces, and that this effect is driven by an increase in neural activity for happy faces. We see this increase in activation for this contrast at both the whole brain level and among our a priori ROIs, specifically the mPFC and bilateral amygdala. Our ROI analyses also indicated increased activation for the intervention condition relative to the control condition in the left amygdala for the happy>sad, happy>fear, happy>sad+fear contrasts. We did not find any differences in neural activation between conditions for our other contrasts at both the whole brain level and in our other ROIs, the bilateral dlPFC and the occipital cortex. Notably, participants in the intervention condition did not show any improvements on measures of depressive symptoms or mood relative to controls following five consecutive days of emotion recognition training.

Our finding of clusters of activation for our happy>sad+fear contrast at the whole brain level in both the left amygdala and brainstem can potentially be explained by the amygdaloid projections underpinning the limbic system. The increase in neural activation for happy expressions for the intervention condition compared to the control condition, resembles changes seen following antidepressant administration. Although effects of SSRIs on amygdala activity in response to positive emotional faces have been reported and replicated, they are less robust than changes in response to negative facial expressions. This is important mechanistically, as anhedonia is characterized by a depressed amygdala response to happy faces (Keedwell et al, 2005)

Increased neural activation to happy faces (and deactivation to sad faces) has been observed following both acute and prolonged antidepressant administration, both in healthy and depressed individuals (Warren, Pringle & Harmer, 2015). Increased activation to positive emotional information following antidepressant treatment has been observed across a large brain network, including the amygdala, mPFC, parahippocampal gyrus, and extra-striate cortex. While these changes may occur in the absence of any effects on participants’ mood, it has been proposed that the early production of a positive bias in emotional processing may be predictive of ultimate symptom improvement in depressed patients (see Warren, Pringle, & Harmer, 2015 for a review). As we did not find any group differences in activation across our contrasts in the bilateral dlPFC and the occipital cortex, we find no evidence that our CBM intervention alters the way an individual attends to emotional expressions, nor does it modify the way these faces are perceived by the visual system. Our analyses suggest that emotion recognition training may increase the salience of positive emotional expressions indexed, by increased neural activation in the amygdala in our intervention v control groups.

While our results indicate that completing a course of emotion recognition training alters neural activation associated with the perception of happy facial expressions, this fMRI study was not powered to detect mood outcomes when comparing participants in intervention and control conditions. Study 2 addresses this question.

## Study Two

### Methods

#### Participants

We recruited adults from the staff and students at the University of Bristol and from the general population who reported depressive symptoms (defined as a score of 14 or more on the BDI-ii). Participants were recruited via email lists and local advertisements.

Upon arrival, participants provided informed consent and inclusion/exclusion criteria were assessed. Screening consisted of the same procedures as in Study One. Criteria for exclusion were also the same as in Study One, except for fMRI-specific contraindications.

The study was approved by the Faculty of Science Research Ethics Committee at the University of Bristol. On completion of the final study session, participants were reimbursed £60 for their time and expenses.

#### Study design and intervention

as in Study 1, participants were allocated to condition using minimisation to balance baseline BDI-ii scores by an experimental collaborator at the Bristol Randomised Trials Collaboration, and testing was double-blind. The CBM intervention and control procedure were the same as in Study One, and participants again completed computerised training or control procedures once a day over five consecutive days (Monday to Friday).

#### Mood Assessment

Mood assessments via questionnaire measures were taken at baseline and at the end-of-treatment. End-of-treatment follow-up included a visual analogue scale rating of how friendly the participant thought the experimenter was, to ensure that there were no differences between treatment conditions that may have affected blinding. The questionnaire measures included the BDI-ii, the BAI, HAM-D, and the PANAS.

#### Other Measures

Friendship and social network size was assessed at baseline by asking participants to rate the number of close friends they currently have on a 5-point scale ranging from 0 (none) to 4 (four or more). A close friend was defined as a person whom respondents report feeling close to and whom they believe they could confide in and turn to for help. Participants repeated this process, rating the number of contacts and acquaintances they currently have. A contact or acquaintance was defined as a person known by sight or known to someone, but not intimately.

Behavioural assessments (Emotion Recognition Task, Scrambled Sentences Test, and the Fishing Game) were taken at the end of treatment, and at 2-week and 6-week follow-up. 6-week follow-up also included a visual analogue scale rating of how helpful the participant thought the experimenter was, to ensure that there are no differences between treatment groups, which may affect blinding. At 6-week follow-up we reassessed the initial exclusion criteria, to establish any change in these.

The emotion recognition task was a 45 trial task that was identical to the baseline block of the training procedure. This was administered to determine whether any chance in bias induced by the task persisted to follow-up. The Fishing Game assessed approach motivation and persistence. In this task, 12 brightly coloured plastic fish move round in a circle, opening and closing their mouths to reveal a magnet (Pictet et al, 2011) Participants are required to catch as many fish as they can in 2.5 minutes by ‘hooking’ them using a magnet on the end of a 900 mm plastic fishing rod. The fishing game task is a simple behavioural performance measure assumed to tap behaviour negatively associated with dysphoria, such as approach motivation and persistence.

The Scrambled Sentences Test assessed depressive interpretation bias. In this task, participants unscramble a list of 20 scrambled sentences (e.g., ‘winner born I am loser a’) under a cognitive load (remembering a six-digit number) (Rude et al 2002)). This task measures the tendency of participants to interpret ambiguous information either positively (‘I am a born winner’) or negatively (‘I am a born loser’). A negativity score is generated by calculating the proportion of sentences completed correctly with a negative emotional valence.

#### Statistical Analysis

We used linear regression to evaluate the effect of training on mood at 6-week follow-up. Analyses were conducted with adjustment for the minimization factor only, and with additional adjustment for age, sex, ethnicity, previous history of treatment for depression, and baseline mood (for analyses of mood variables only). The primary outcome was depressive symptoms over the last 2 weeks assessed using the BDI-ii at 6-week follow-up. Secondary outcomes included depressive symptoms measured using the HAM-D, and positive and negative affect assessed using the PANAS. Subgroup analyses were conducted stratified by whether participants meet criteria for clinical depression, number of episodes of depression, age at first episode, and whether participants had depression with or without anxiety. We also analysed the impact of social network size on treatment effect.

Our preliminary data indicated an effect size of *d* = 0.43 at 2-week follow-up, corresponding to a difference of 3 points on the Positive and Negative Affect Schedule (PANAS). This suggested that a sample size of 172 would be required to achieve 80% power at an alpha level of 5%. This sample size gave us equivalent power to detect a difference of 5 points on the BDI-ii at 6-week follow-up (our primary outcome), which we considered would be clinically significant. We aimed to recruit 190 participants to accommodate potential attrition. The study protocol was registered prior to data collection (ISRCTN17767674) (Adams et al, 2013). All analyses were conducted using SPSS Statistics version 21.

### Results

#### Characteristics of Participants

A total of 190 participants (138 female) aged 18 to 39 years (*M* = 21, *SD* = 4) were recruited. The characteristics of participants by condition are shown in Table 1.

#### Primary Outcome

We found no clear evidence of a reduction in depressive symptoms on the BDI-ii at 6-week follow-up (our primary outcome) in the intervention condition compared with the control condition in either the unadjusted (mean difference 0.35, 95% CI −2.41 to 3.10, P = 0.80) or adjusted (mean difference 0.10, 95% CI −2.39 to 2.58, P = 0.94) models.

#### Secondary Outcomes

There was also no clear evidence of a difference between the two conditions on the BDI-ii at any other time points, or on any other mood measures. These results are shown in Table 3. We also found no clear evidence of a difference on the Scrambled Sentences Test (unadjusted mean difference 0.48, 95% CI −0.94 to 1.90, P = 0.51; adjusted mean difference 0.30, 95% CI −1.12 to 1.72, P = 0.68), or the Fishing Game (unadjusted mean difference 0.23, 95% CI −2.24 to 2.70, P = 0.85; adjusted mean difference 0.28, 95% CI −2.24 to 2.79, P = 0.83) at 6-week follow-up. However, we did find clear evidence of a difference on the Emotion Recognition Task at 6-week follow-up (unadjusted mean difference −2.91, 95% CI −3.67 to −2.14, P < 0.001; adjusted mean difference −2.84; 95% CI −3.63 to −2.06, P < 0.001), indicating that the effect of the intervention on cognitive bias persisted beyond the treatment phase. Similar results were obtained for these measures at end-of-treatment and 2-week follow-up.

**Table 3.** **Effects of Emotion Recognition Training on Mood Symptoms in Study 2.**

|  |  | End of Treatment |  |  | Follow-Up (2 weeks) |  |  | Follow-Up (6 weeks) |  |  |
| --- | --- | --- | --- | --- | --- | --- | --- | --- | --- | --- |
|  |  | Estimate | 95% CI | P | Estimate | 95% CI | P | Estimate | 95% CI | P |
| BDI-II | Unadjusted | -0.59 | -3.33 to 2.15 | 0.67 | -0.26 | -3.18 to 2.66 | 0.86 | 0.35 | -2.41 to 3.10 | 0.80 |
|  | Adjusted | -0.50 | -2.50 to 1.50 | 0.62 | -0.75 | -3.11 to 1.60 | 0.53 | 0.10 | -2.39 to 2.58 | 0.94 |
| BAI | Unadjusted | -0.19 | -2.50 to 2.12 | 0.87 | -0.34 | -3.08 to 2.39 | 0.81 | -0.15 | -2.77 to 2.47 | 0.91 |
|  | Adjusted | -1.14 | -2.56 to 0.28 | 0.12 | -1.71 | -3.96 to 0.53 | 0.13 | -1.10 | -3.36 to 1.16 | 0.34 |
| HAM-D | Unadjusted | -0.21 | -1.72 to 1.30 | 0.79 | 0.17 | -1.61 to 1.94 | 0.85 | 1.02 | -0.62 to 2.65 | 0.22 |
|  | Adjusted | -0.36 | -1.52 to 0.81 | 0.55 | -0.18 | -1.56 to 1.20 | 0.80 | 0.80 | -0.63 to 2.24 | 0.27 |
| PANAS Positive | Unadjusted | 0.88 | -1.14 to 2.89 | 0.39 | 1.35 | -0.80 to 3.49 | 0.22 | 0.17 | -2.20 to 2.54 | 0.89 |
|  | Adjusted | 0.20 | -1.37 to 1.76 | 0.81 | 0.49 | -1.31 to 2.28 | 0.59 | -0.44 | -2.54 to 1.66 | 0.68 |
| PANAS Negative | Unadjusted | -0.52 | -1.96 to 0.91 | 0.47 | 0.49 | -1.17 to 2.14 | 0.56 | -0.02 | -1.43 to 1.38 | 0.97 |
|  | Adjusted | -0.63 | -1.70 to 0.45 | 0.25 | 0.36 | -0.95 to 1.67 | 0.59 | -0.05 | -1.35 to 1.25 | 0.94 |

BDI-ii: Beck Depression Inventory; HAM-D Hamilton Rating Scale for Depression; PANAS: Positive and Negative Affect Schedule. Adjusted analyses include age, sex, ethnicity, previous history of treatment for depression, and baseline mood score as covariates.

#### Planned Sub-Group Analyses

Subgroup analyses, both unadjusted and adjusted, did not indicate any clear evidence of improved mood in the intervention condition compared to the control condition among participants with a diagnosis of clinical depression, number of previous episodes of depression, age at first episode among those with a previous episode, and among those with high levels of anxiety symptoms.

#### Unplanned Exploratory Analyses

Given the lack of an effect of CBM on mood, we explored whether emotion recognition bias may instead serve as a cognitive biomarker for depressed mood, by calculating the correlation between pre-training balance point at session 1 and self-reported measures of mood at the same time point. We found evidence of consistent, albeit relatively weak, correlations across most measures (BDI-ii: r = −.18, P = .018; HAM-D: r = −.17, P = .021; BAI: r = −.11, P = .12; PANAS Positive: r = +.23, P = .001; PANAS Negative: r = -.03, P = .67).

### Discussion

Our results suggest that emotion recognition training induces a change in cognitive bias that persists for 6 weeks after the end of treatment but does not reduce depressive symptoms on the BDI-ii, or on related measures of mood, motivation and persistence, or depressive interpretation bias. We did not find any evidence of specific sub-groups that benefited from the intervention in our planned sub-group analyses. However, we did find evidence that emotion recognition bias may serve as a cognitive biomarker for depressed mood (and in particular low positive affect).

## General Discussion

These two studies present evidence that a simple, automated CBM task leads to training effects that increase amygdala response to happy faces (Study 1) and persist for at least six weeks (Study 2). There is no evidence, however, that this cognitive bias modification has any effects on either questionnaire measures of mood, or behavioural measures of anhedonia (the fishing task). Given the robust nature of the training effects, these findings question theories that suggest a strong causal role for emotion processing biases in the onset or maintenance of depression.

One possibility is that the emotional training task does not generalize to other situations in which any therapeutic effects of modified bias may be realized (e.g., social interactions). Although we know that the training effect transfers to other faces in an experimental context (e.g., the face task in Study 1, which employs different faces to the training task, see also Dalili et al, 2017), there is currently little evidence that this bias generalizes to real world encounters with others. A recent study of a similar bias modification task delivered online (Peters et al, 2017) to healthy participants showed little evidence of transfer of bias modification to a variety of cognitive tasks thought to be impacted by low mood (although there was weak evidence of transfer to measures of stress and a cognitive measure of anhedonia, particularly in those participants with higher baseline anxiety). A further RCT employing a modified version of the CBM technique reported here aiming to reduce social anxiety in adolescent participants also showed weak results. Although there was no decrease in social anxiety (the primary outcome), participants in the intervention group showed lower depression symptoms at 2-week follow up (Fitzgerald et al 2018).

Alternatively, individual differences in emotion processing may play no causal role in the onset or maintenance of depression. Instead, emotion processing could be a useful cognitive biomarker of depressive mood, and may act as a marker of treatment success.

## Acknowledgements

MRM and SA are members of the UK Centre for Tobacco and Alcohol Studies, a UKCRC Public Health Research: Centre of Excellence. Funding from British Heart Foundation, Cancer Research UK, Economic and Social Research Council, Medical Research Council, and the National Institute for Health Research, under the auspices of the UK Clinical Research Collaboration, is gratefully acknowledged. This work was supported by the Medical Research Council (MR/J011819/1).

